# Large Serine Integrase Off-target Discovery and Validation for Therapeutic Genome Editing

**DOI:** 10.1101/2024.08.23.609471

**Authors:** Dane Z. Hazelbaker, Japan B. Mehta, Connor McGinnis, Didac Santesmasses, Anne M. Bara, Xiaoyu Liang, Thomas Biondi, Tim Fennell, Nils Homer, Brett Estes, Jenny Xie, Davood Norouzi, Kaivalya Molugu, Ravindra Amunugama, Chong Luo, Parth Amin, Xiarong Shi, Jesse Cochrane, Sandeep Kumar, Jie Wang, Matthew H. Bakalar, Jonathan D. Finn, Daniel J. O’Connell

## Abstract

While numerous technologies for the characterization of potential off-target editing by CRISPR/Cas9 have been described, the development of new technologies and analytical methods for off-target recombination by Large Serine Integrases (LSIs) are required to advance the application of LSIs for therapeutic gene integration. Here we describe a suite of off-target recombination discovery technologies and a hybrid capture validation approach as a comprehensive framework for off-target characterization of LSIs. HIDE- Seq (High-throughput Integrase-mediated DNA Event Sequencing) is a PCR-free unbiased genome-wide biochemical assay capable of discovering sites with LSI-mediated free DNA ends (FDEs) and off-target recombination events. Cryptic-Seq is a PCR-based unbiased genome-wide biochemical or cellular-based assay that is more sensitive than HIDE-Seq but is limited to the discovery of sites with off-target recombination. HIDE-Seq and Cryptic-Seq discovered 38 and 44,311 potential off-target sites respectively. 2,455 sites were prioritized for validation by hybrid capture NGS in LSI- edited K562 cells and off-target integration was detected at 52 of the sites. We benchmarked the sensitivity of our LSI off-target characterization framework against unbiased whole genome sequencing (WGS) on LSI-edited samples, and off-target integration was detected at 5 sites with an average genome coverage of 40x. This reflects a greater than 10-fold increase in sensitivity for off-target detection compared to WGS, however only 4 of the 5 sites detected by WGS were also validated by hybrid capture NGS. The dissemination of these technologies will help advance the application of LSIs in therapeutic genome editing by establishing methods and benchmarks for the sensitivity of off-target detection.

## INTRODUCTION

The large serine integrase (LSI) family constitutes a diverse group of site-specific recombinases that play pivotal roles in mediating DNA rearrangements^1-3^. Serine integrases, in contrast to their tyrosine recombinase counterparts, utilize a serine residue for catalysis, leading to distinct mechanistic features^4^. This large family encompasses integrases with varying sizes and functionalities, with notable members including PhiC31 integrase from *Streptomyces* bacteriophage PhiC31^5^ and Bxb1 integrase discovered in mycobacteriophage Bxb1^6^. Both PhiC31 and Bxb1 integrases are well-recognized for their utility in site-specific recombination applications by virtue of direct recombination between phage attachment site *attP* and bacterial attachment site *attB* with the requirement of no co-factors or DNA supercoiling^7-9^. The precise and efficient DNA manipulation capabilities of the large serine integrase family have positioned it as an attractive tool for synthetic biology and genome editing applications^10-14^.

Large serine integrases facilitate recombination between attachment sites on linear or circular DNA substrates^5, 15^ and the mechanism of action is outlined in **Figure 1a**. Dimerized integrases bind to specific sequences in phage (*attP*) and bacterial host (*attB*) DNA^2^ The bound integrases associate to form a synaptic complex, connecting paired homologous sequences. Integrases cleave all four DNA strands at the central dinucleotide, forming 5′-phosphoserine linkages and generating 3′-dinucleotide overhangs^3^. This cleavage mechanism covalently bonds DNA strands to integrase, avoiding free DNA ends and the opportunity for mis-repair by host DNA machinery. Subunits exchange places, promoting irreversible ligation through the formation of new attachment sites, *attL* and *attR*. Unlike CRISPR/Cas9 knock-in approaches that depend on cellular DNA repair machineries^5, 15^, LSI integration is deterministic and independent of host cell factors ^5, 15^.

**Figure 1.**
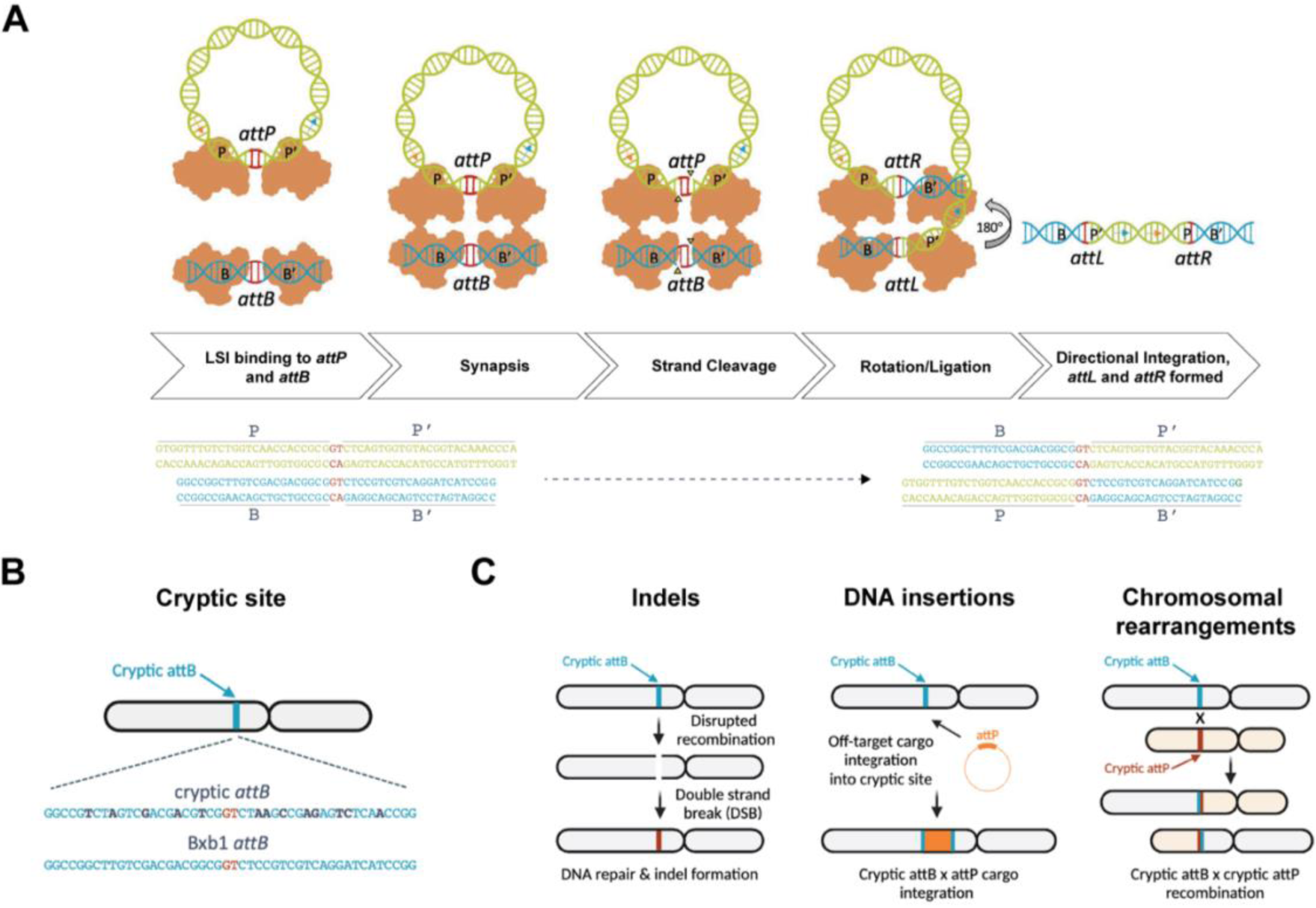
Large Serine Integrase (LSI) Mechanism of Action. **(A)** Serine integrases catalyze recombination between attachment (*att*) sites on linear or circular DNA substrates. Integrase dimers bind to specific att sequences in the phage (*attP*) and bacterial host (*attB*) DNA. Integrase bound to *attP* and integrase bound to *attB* associate to form a synaptic complex that connects the paired homologous sequences. The integrase subunits cleave all four DNA strands at the central dinucleotide, forming 5′-phosphoserine linkages between integrase subunits and DNA half-sites and generating 3′-dinucleotide overhangs. The P′ and B′-linked subunits exchange places by rotating 180° about a horizontal axis relative to the P and B-linked subunits. Base-pairing between the central dinucleotides promotes ligation of the DNA strands, resulting in formation of two new attachment sites, attachment left (*attL*) and attachment right (*attR*). **(B)** Loci in the human genome with homology to LSI *attB* or *attP* sites are often called cryptic attachment sites because they are not obvious but have sufficient homology to catalyze recombination. **(C)** There are 2 classes of off-target editing from an LSI; DNA mutagenesis in the form of indels from FDEs that may occur if recombination is disrupted in between strand cleavage and re-ligation. The second class is DNA structural variants, and these can come in two forms, the first is from off-target cargo insertion at a site in the human genome with homology to the attachment sequences. The second appears in the form of gross chromosomal rearrangements that may involve the attachment sequence introduced by genome editing or the interaction between two sites with homology to the attachment sequences.

A comprehensive understanding of the specificity of integrase-mediated insertion for gene editing strategies, such as integrase-mediated programmable genomic integration (I-PGI), is crucial for safe and effective therapeutic use. While LSIs attachment sites of *attP* and *attB* are typically between 40-50 bp^16^ and recombine efficiently with a high degree of site-specificity^2^, sites with degenerate homology to *attB* or *attP*, known as cryptic sites or pseudo sites can exist in the human genome and can support recombination at lower frequencies^12, 17, 18^ (**Figure 1b**). In terms of off-target editing events potentially mediated by cryptic sites in I-PGI, and other integrase-based editing approaches^12, 13^, we broadly classify these events into three genetic classes: indels, DNA insertions, and large chromosomal rearrangements (**Figure 1c**). In the case of indels, because site-specific recombination by LSIs proceeds through a cleaved DNA intermediate, disruption of this intermediate could result in de-protection of the cleaved free DNA ends (FDEs) and repair of the exposed DNA ends by the cellular DNA repair^19^. Biochemical evidence for FDE formation via disruption of LSI recombination by chemical treatment or mismatched central dinucleotides exists in biochemical systems^15^. In the case of off-target integration events, Bxb1 integrase has been reported to integrate DNA cargoes into a small number of sites in the human genome at low levels^12, 18^, though the extent of potential Bxb1 integrase cryptic sites in the human genome for Bxb1 is not clear. In addition to off-target recombination of DNA cargo into cryptic sites, there exists the possibility of recombination between two cryptic sites in the human genome with previous publications describing multiple chromosomal rearrangements by PhiC31 integrases in human cells, potentially mediated by cryptic sites in the human genome^20^ (**Figure 1c**). To address these potential outcomes, we set out to develop more sensitive methods to determine the off-target profile of Bxb1 integrase in human cells.

Here we demonstrate an empirical off-target discovery and validation framework for the characterization of LSI specificity for therapeutic genome editing that is sensitive and comprehensive of unintended DNA variants. We describe the generation of a suite of sensitive biochemical-based discovery and characterization approaches in addition to a targeted hybrid capture NGS approach for the validation of off-target LSI editing in cells at sites nominated by our discovery assays. For discovery of genomic sites that support LSI off-target activity, we present HIDE-seq and Cryptic-seq. HIDE-Seq (High-throughput Integrase-mediated DNA Event Sequencing) is a PCR-free unbiased genome-wide biochemical assay capable of discovering sites with LSI-mediated FDEs and off-target recombination events. Cryptic-Seq is a PCR-based unbiased genome-wide biochemical or cellular-based assay that is more sensitive than HIDE-Seq but is limited to the discovery of sites with off-target recombination. The application of these biochemical characterization technologies in combination with cell-based hybrid capture NGS validation, help advance the implementation of LSIs in therapeutic genome editing approaches by establishing benchmark criteria for sensitive off-target discovery and validation in therapeutically relevant cell-types.

## RESULTS

### HIDE-Seq is an unbiased and genome-wide LSI off-target discovery technology

For unbiased and genome-wide assessment of potential LSI off-target editing events, we combined a biochemical LSI recombination reaction containing recombinant integrase, substrates and purified and deproteinized human genomic DNA with whole genome sequencing. We named this approach HIDE-seq (<u>H</u>igh-throughput Integrase-mediated <u>D</u>NA <u>E</u>vent Sequencing), a PCR-free genome-wide biochemical assay for discovering LSI-mediated off-target events, including FDEs, cryptic integration events, or structural variants via WGS (**Figure 2a**). For the development of HIDE-seq, we drew inspiration from Digenome-seq^21^, a biochemical assay for the genome-wide detection of CRISPR-Cas9 off-target cleavage via WGS of DNA samples digested with Cas9 to identify read pairs reads that correspond to FDEs.

**Figure 2.**
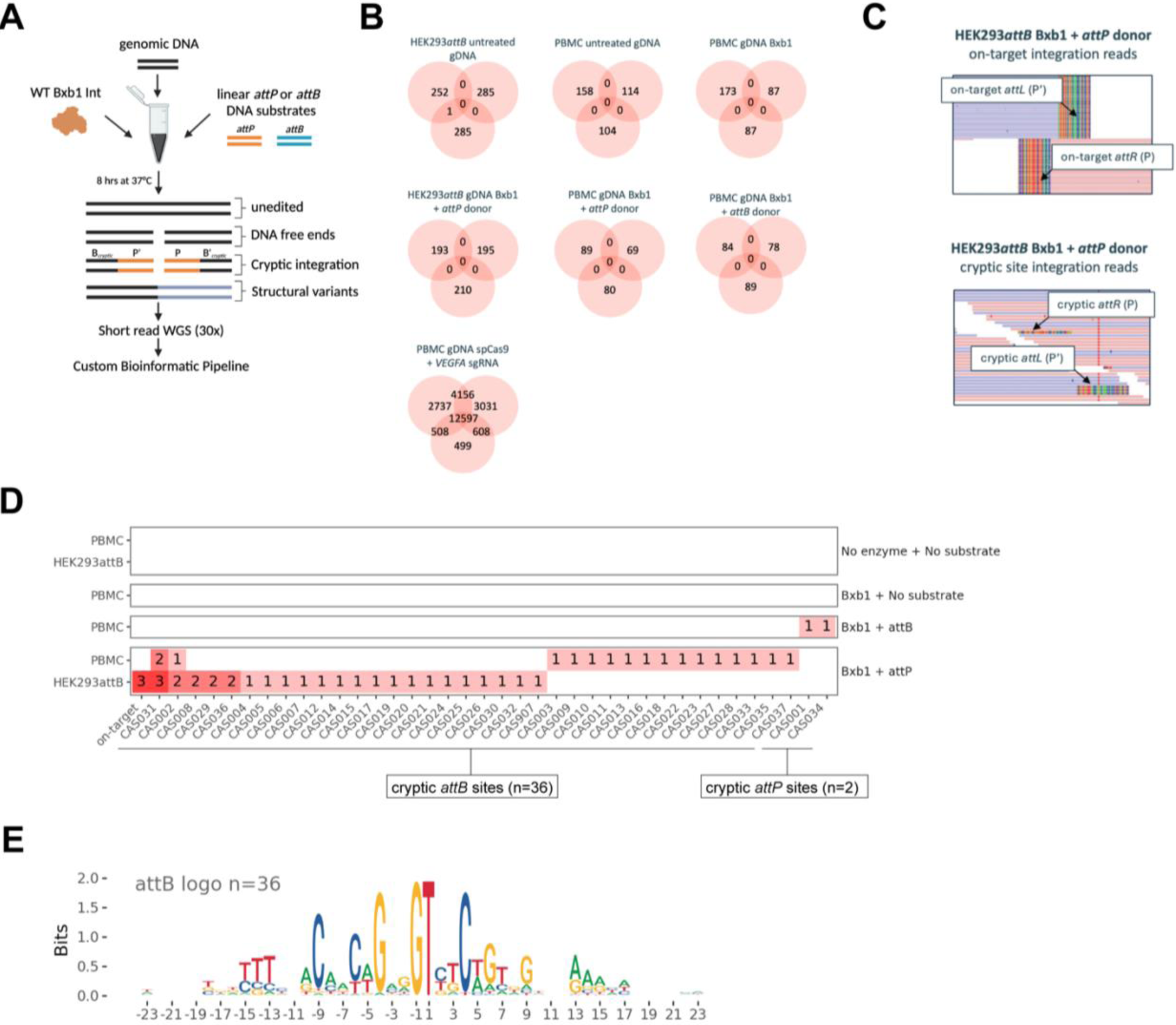
High-throughput Integrase-mediated DNA Event Sequencing (HIDE-seq) **(A)** Graphic of HIDE-seq experimental workflow. Genomic DNA is recombined with linear *attB* or *attP* substrates by an LSI and then subjected to WGS **(B)** Venn diagrams of the loci from FDEs detected by HIDE-seq for Bxb1 and Digenome-seq for Cas9. Each circle represents a biochemical replicate reaction. Zero recurrent and background levels of FDEs were detected in samples treated with Bxb1 alone, Bxb1 and *attP*, or Bxb1 and *attB*. In contrast, 12,597 recurrent FDEs were detected in our in-line control using Digenome-seq and Cas9 with the off-target standard sgRNA VEGFA site 2. **(C)** Integrative Genomics Viewer (IGV) of HIDE-seq reads supporting recombination display distinct opposing read alignments. The top panel corresponds to on-target *attB* integration reads supporting recombination with *attL* (P’) and *attR* (P) sequences appended to the ends. The reads align with lentiviral genomic sequence in the HEK293*attB* genome and then appear soft-clipped (multicolored) where the linear *attP* substrates were recombined because these sequences do not exist in the human genome hg38 alignment reference file. The bottom panel corresponds to off-target integration where soft clipped reads that correspond to *attL* (P’) and *attR* (P) sequences are detected amongst the WGS coverage. In contrast to the on-target locus where all reads covering the locus were recombined, only a fraction of reads appear recombined at potential off-target loci. **(D)** HIDE-seq discovered 36 cryptic *attB* and 2 cryptic *attP* loci in the human genome. **(E)** DNA sequence motif logo of cryptic *attB* sites created by aligning the 46 bp sequence from the 36 cryptic *attB* sites.

To demonstrate the utility of HIDE-seq, we leveraged the well-characterized LSI Bxb1 integrase^6, 22^ and isolated human genomic DNA (gDNA) from two sources (1) a HEK293 cell line that contains two copies of the Bxb1 *attB* site in a lentiviral vector integrated into a single locus on chromosome 5 (referred to from here on as HEK293*attB*) and (2) commercially available peripheral blood mononuclear cells (PBMCs).

The cell-free nature of HIDE-seq allows for tuning and supraphysiological saturation of LSI and DNA substrates in the reaction to enable sensitive discovery of potential off-target events to deeply map the landscape of potential off-target events across any given DNA sample. For HIDE-seq reactions, 8 μg of PBMC or HEK293*attB* gDNA was incubated with 10 nM *attP* or *attB* linear substrate and 400 nM of recombinant Bxb1 integrase in various reactions in triplicate for 8 hours at 37C° as outlined in **Table 1**. As a positive control reaction for free DNA end formation, we performed Digenome-seq^21^ reactions with Cas9 complexed with a previously described low specificity sgRNA^23^ (**Table 1**). After completion of the reactions, the gDNA was purified and processed via amplification-free library preparation and subjected to Illumina WGS with a target of 30x coverage (recovered mean coverage of 18.5x, with a minimum of 10x and maximum of 28x across samples) followed by bioinformatics analysis by Digenomitas^24^ (for FDE discovery) or a custom HIDE-seq pipeline (for LSI off-target discovery).

**Table 1.** HIDE-seq sample summary. Overview of HIDE-seq samples performed in triplicate (3x) and the experimental purpose.

| Sample (x3) | Purpose |
| --- | --- |
| PBMC untreated gDNA | PBMC unedited control DNA sample |
| PBMC gDNA spCas9 +*VEGFA* sgRNA | PBMC FDE positive control sample |
| PBMC gDNA Bxb1 | PBMC Bxb1 events |
| PBMC gDNA Bxb1 + *attB* donor | PBMC Bxb1 and *attB* donor events |
| PBMC gDNA Bxb1 + *attP* donor | PBMC Bxb1 and *attP* donor events |
| HEK293*attB* untreated DNA | HEK293*attB* unedited control DNA sample |
| HEK293*attB* Bxb1 + *attP* donor | HEK293*attB* Bxb1 and *attB* donor events |

No LSI associated FDE’s were identified, as Bxb1 treated samples had a similar number of FDEs (69-212 total counts) to background levels of untreated DNA samples (104-206 total counts) (**Figure 2b**). Furthermore, no recurrent FDEs were detected by HIDE-seq, where recurrence is defined as the appearance of a FDE at the same genomic location in the triplicate samples (**Figure 2b**). In contrast, 12,597 recurrent FDEs were detected in samples treated with spCas9 complexed with VEGFA sgRNA (**Figure 2b**). These results indicate that Bxb1 integrase does not generate significant levels of FDEs during its canonical recombination mechanism consistent with previous results shown for Bxb1 in biochemical contexts^15^.

HIDE-seq identified high levels of on-target integration between the *attP* donor and the endogenous *attB* sites in the HEK293*attB* gDNA samples, with integration read count frequencies of 100%, 100%, and 94% for HEK293*attB* Bxb1 + *attP* donor replicates 1, 2, and 3 respectively (quantified as percentage of *attL* and *attR* reads versus *attB* total read counts, data not shown). An example visualization image of HEK293*attB* Bxb1 + *attP* donor replicate 1 displaying 100% integration of the *attP* donor in integrative genomics viewer^25^ (IGV) is shown in **Figure 2c**. These high levels of on-target integration indicate our biochemical reaction conditions and concentrations of Bxb1 integrase and DNA donor substrates are robust enough to detect potential off-target events, as shown by the integration reads at an example cryptic *attB* site (**Figure 2c)**. Across the aggregate of PBMC and HEK293*attB* gDNA samples treated with Bxb1 and *attP* donor, we discovered 36 unique cryptic *attB* sites, which we designate as CAS sites (<u>c</u>ryptic <u>a</u>ttachment <u>s</u>ites) (**Figure 2d and Supplementary Table 1**). For the PBMC gDNA samples treated with Bxb1 and *attB* donor, we discovered 2 unique cryptic *attP* sites. Our detection of fewer numbers of cryptic *attP* sites compared to cryptic *attB* sites is consistent with recently published results^18^. With the 36 cryptic *attB* sites uncovered by HIDE-seq, we generated detailed sequence logos that represent the specificity profile of Bxb1 across the human genome (**Figure 2e**). Lastly, to examine if any split reads in the HIDE-seq dataset support potential recombination between any of the 38 discovered cryptic sites to another genomic location, we utilized fgsv caller by Fulcrum Genomics^26^. With fgsv caller analysis, we found no split reads that map at both the cryptic site and another genomic location, suggesting that these 38 discovered cryptic sites did not engage in detectable cryptic recombination in the HIDE-seq reactions (data not shown). Taken together, these results demonstrate HIDE-seq is a sensitive and unbiased approach to identify any potential FDEs or off-target integration events by LSIs.

### Cryptic-Seq is a sensitive discovery technology for LSI off-target recombination

While HIDE-seq can identify potential LSI cryptic sites via sequencing whole genomes, we sought to develop a genome-wide approach that preferentially enriches recombined species to enable more sensitive discovery of cryptic sites. We named this approach Cryptic-seq, and it relies on the application of a specialized DNA donor substrate to enable one-step PCR enrichment and sequencing of recombined cryptic sites (**Figure 3a**). To execute Cryptic-seq, gDNA is first tagmented with Tn5 transposase^21^ loaded DNA oligonucleotides containing Illumina P5 sequencing adaptors, i5 indexes, and a unique molecule identifier (UMI)^27-29^ (**Figure 3a**). This pre-tagmentation of the gDNA is a crucial step to increase the sensitivity of the assay as it prevents sequencing of any unrecombined donor DNA substrate. After tagmentation, gDNA is incubated with recombinant Bxb1 integrase and a Cryptic-seq plasmid-based vector that contains either *attP* (GT dinucleotide) flanked by a unique barcode sequence (BC1) and an Illumina N7 primer binding site 5’ of the P half-site of *attP,* and on 3’ side of the P’ site of the *attP* a second unique barcode sequence (BC2) and an Illumina P7 primer binding site (**Figure 3a**). After the reaction is complete, the gDNA is purified and PCR amplified with i7 indexed primers that prime off either the N7 or P7 primer binding site in the cryptic-seq vector for one-step amplification of recombined DNA fragments (**Figure 3a**), followed by NGS and bioinformatic identification of genomic reads sequence-tagged with either the P or P’ half-sites from the *attP* donor. Like HIDE-seq, Cryptic-seq is a biochemical-based assay and the reactions are performed with supraphysiological concentrations of Bxb1 integrase and DNA donor, conditions far higher than we can achieve in a cell, for maximum sensitivity to enable deep discovery of any potential off-target events.

**Figure 3.**
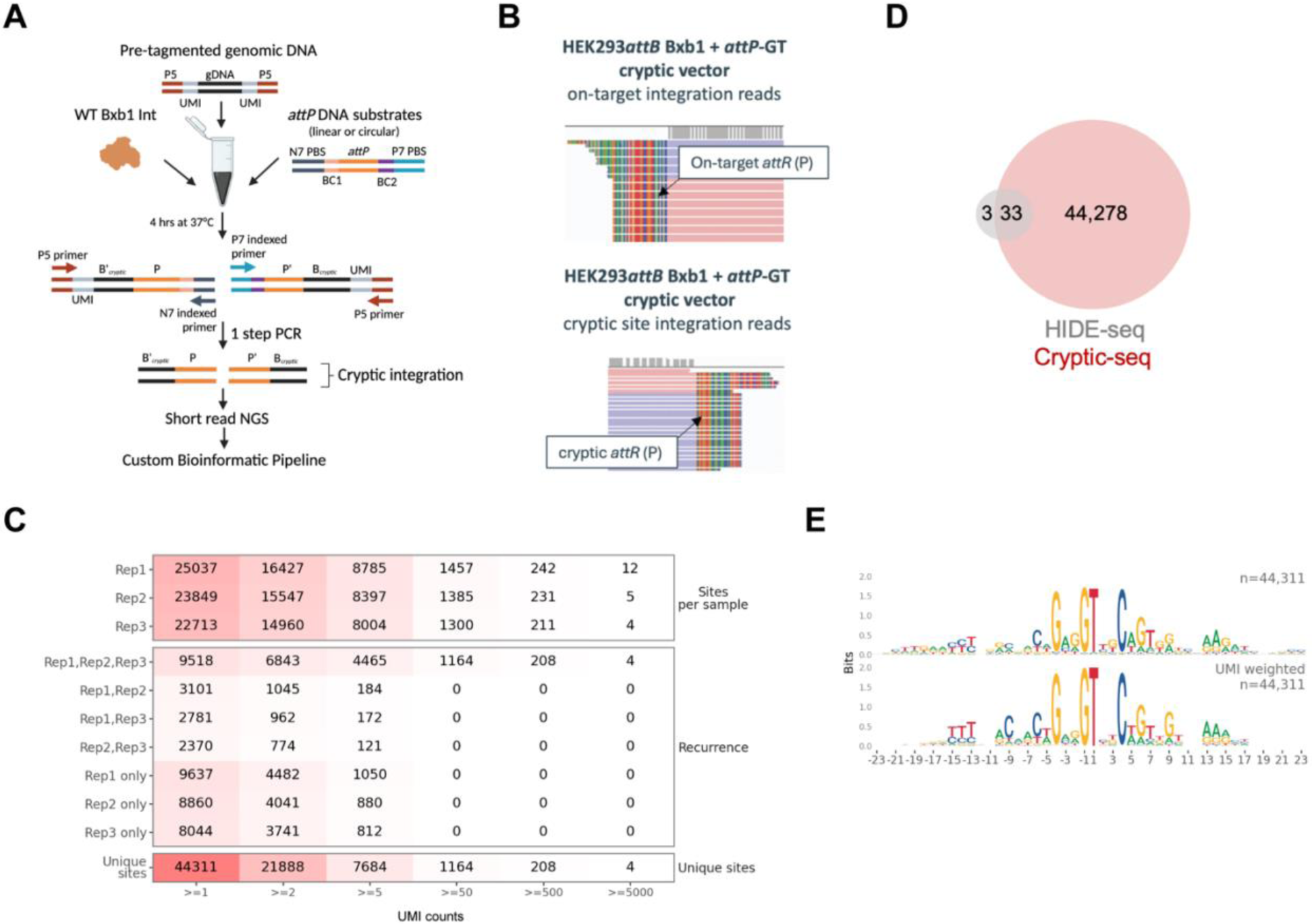
Cryptic-seq. **(A)** Graphic of Cryptic-seq experimental workflow. First, a library of genomic DNA is created with Tn5 tagmentation that imparts partial Illumina sequencing adapters. This library of genomic DNA is then recombined with an LSI and *attP* or *attB* DNA substrates with complementary Illumina sequencing adapters. Potential off-target loci in the human genome that are recombined by the LSI are subject to one-step PCR enrichment and deep sequencing. **(B)** Representative IGV browser of Cryptic-Seq reads supporting recombination at the on-target *attB* site and a cryptic *attB*. The top panel corresponds to on-target alignment of the left library (N7) with reads supporting the right junction (*attR*). The reads that align with lentiviral genomic sequence in the HEK293*attB* genome appear soft-clipped where the linear *attP* substrates were recombined because these sequences do not exist in the human genome hg38 alignment reference file. The bottom panel corresponds to off-target alignment of the N7 library with reads supporting *attR* (in this view the cryptic site is in the antiparallel orientation such that the GT dinucleotide is on the bottom DNA strand so the *attR* reads appear on the left). The reads align with genomic sequence and then appear soft-clipped where the linear *attP* substrates were recombined because these sequences do not exist in the human genome hg38 alignment reference file. **(C)** Summary table with the number of cryptic attB sites detected in the human genome. The “Sites per sample” section at the top indicates the total number of sites detected in each replicate, at increasing UMI detection level, up to >= 5000 unique recombination events. The “Recurrence” section summarizes the overlap analysis across replicates. The number of sites shared between the corresponding set of replicates is indicated, e.g., the top row “Rep1,Rep2,Rep3” corresponds to sites observed in all three Cryptic-seq replicates, at increasing UMI levels. For sites shared between replicates, the lowest of the UMI values is used. As shown here, all sites ≥ 50 UMIs were observed in all three replicates. The bottom row “Unique sites” section indicates the total number of unique sites across all three replicates, at increasing UMI levels. A total of 44,311 unique cryptic *attB* sites were detected in the hg38 reference human genome with 1 UMI or more. **(D)** Venn diagram of cryptic *attB* discovery overlap from HIDE-Seq and Cryptic-seq. **(E)** DNA sequence motif logos, unweighted by UMI count (top) and weighted by UMI count (bottom), created by aligning the 46 bp sequence from the 44,311 cryptic *attB* sites.

To apply Cryptic-seq to the human genome, we isolated gDNA from HEK293*attB* cells and used Tn5 transposase to tagment the library to an average fragment size of 700 bp. Three independent replicate reactions were performed at 37C° for 4 hours with each reaction containing 1 µg tagmented HEK293*attB* gDNA, 10 nM of the attP-GT vector, and 1 µM Bxb1 integrase. After quenching of the reaction with sodium dodecyl sulfate (SDS) the gDNA was purified and PCR enrichment was performed with N7 primer. In this report, Cryptic-seq was performed with one-sided enrichment, with only indexed N7 primers used for detection of cryptic *attR* fragments because the creation of *attL* from a cryptic *attB* is inextricably linked to the formation of *attR* by Bxb1 integrase. After PCR-based enrichment and indexing, NGS libraries were quantified and sequenced by Illumina next generation sequencing on a NextSeq 2000. Examples of representative IGV^25^ plots showing the structure of both on-target and cryptic site genomic reads tagged with the P half-site are shown in **Figure 3b**.

In total across the three replicates (Cryptic-seq Rep1, Rep2, and Rep3), 44,311 unique cryptic *attB* sites with ≥ 1 UMI support were discovered by Cryptic-seq, with each replicate containing > 22,000 discovered cryptic sites in the human genome that can recombine in a Bxb1 integrase-dependent manner (**Figure 3c and Supplementary Table 2**). The recombination signal produced by Cryptic-Seq is reflected by the number of UMIs detected for each site and the distribution is negatively skewed with a long-tail of single UMI events detected. In each replicate, over 200 sites show greater than 500 UMIs counted, reflecting that these sites have recombined individual and unique DNA fragments containing these cryptic sites over 500 times in the Cryptic-seq reaction (**Figure 3c)**. Of note, 4 unique cryptic sites displayed UMI counts greater than 5000 UMIs in all three replicates (**Figure 3c**). For reference, the endogenous *attB* site in the HEK293*attB* genome displayed integration read UMI counts of 6033 UMIs in Rep1, 5622 UMIs in Rep2, and 4808 UMIs in Rep3, indicative of high levels of on-target recombination (data not shown). The recurrence of recombination at these sites varies also across replicates, with sites with high UMI counts (≥ 50 UMIs) displaying higher levels of recurrence than sites with low UMI counts (≤ 5 UMIs) (**Figure 3c**).

Cryptic-Seq re-discovered 33 of the 36 cryptic *attB* sites that were discovered by HIDE-seq (**Figure 3d**), which reflected a 91% concordance in that comparison. However, Cryptic-seq also discovered an additional 44,278 cryptic *attB* sites in the human genome. Despite this large increase in the number of loci detected, the Bxb1 sequence motif (**Figure 3e**) did not change significantly compared to the HIDE-seq data, which indicates that these data represent a 1,265-fold increase in sensitivity and not the result of spurious biochemical signals.

### Validation of Hybrid Capture NGS as a scalable approach for LSI editing quantification

With a >1200-fold increase in sensitivity for LSI cryptic site discovery by Cryptic-seq over HIDE-seq and thousands of potential off-target sites discovered, we required a scalable validation approach to characterize off-target editing frequencies in a relevant cellular context. Droplet digital PCR^30^ (ddPCR), while quantitatively accurate, cannot be readily scaled to thousands of unique sites. Multiplex targeted NGS amplicon sequencing assays, such as rhAmpSeq can scale to thousands of sites for approaches like CRISPR/Cas9 editing detection^31^, however the unique sequence of recombined reads generated by LSIs does not allow for quantitative detection with forward and reverse genomic primers surrounding the target site. To overcome these limitations, we turned to multiplex hybrid capture NGS^32^ which has been applied previously to assess off-target editing frequencies of thousands of potential off-target sites of the first FDA approved CRISPR-based therapeutic, Exagamglogene autotemcel (marketed as CASGEVY)^33, 34^

To customize hybrid capture NGS for the detection of LSI-editing events (indels, cryptic integration events, and structural variants) in DNA from edited cells, we first designed 120-mer DNA capture probes to specifically bind left (5’) and right (3’) of the discovered cryptic site central dinucleotide (**Figure 4a**). By designing probes flanking both sides of the central dinucleotide of the cryptic site, we could compare the capture and quantification efficiency of the left and right probes. Importantly, by designing probes that flanking the central dinucleotide of the genomic site, we ensure the quantitative nature of the assay is preserved as the capture probe will enrich both edited and unedited genomic sites with equal efficiency, thus allowing accurate quantification of edited and unedited DNA species after bioinformatic UMI deduplication of NGS reads.

**Figure 4.**
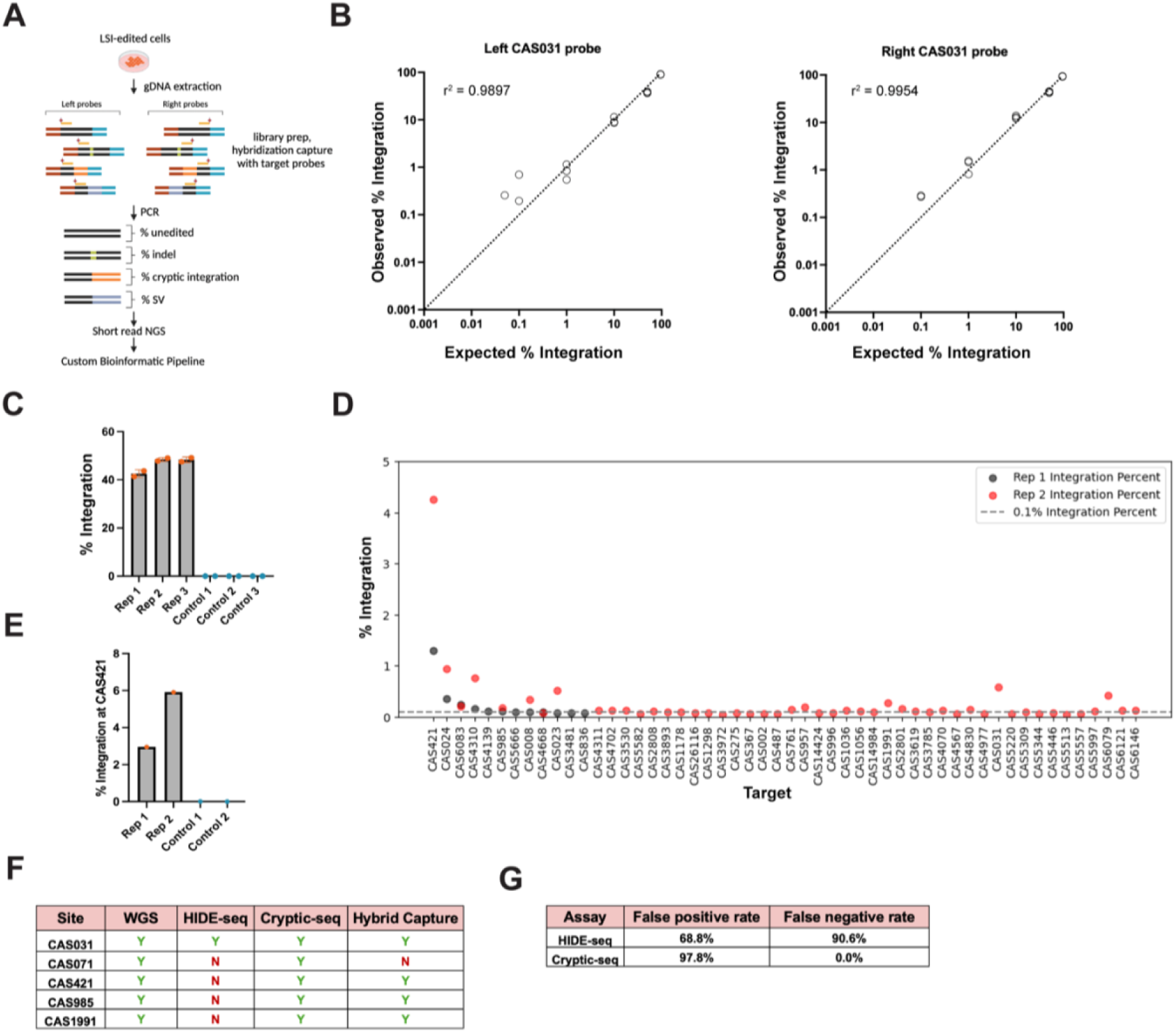
Hybrid Capture NGS Validation of Off-Target Editing. **(A)** K562*attB* cells were edited with Bxb1 integrase and the purified gDNA was used for hybrid capture followed by NGS. Hybrid capture probes were designed on the left and right sides of the cryptic attachment loci to enable unbiased detection of indels, cargo integration and genomic rearrangements. **(B)** The sensitivity of off-target insertion detection with hybrid capture NGS was 0.1% when using a synthetic DNA standard spike-in from one of the off-target loci discovered, CAS031. r^2^= Pearson coefficient of determination **(C)** K562*attB* cells edited with Bxb1 had 45-48% levels of insertion measured by ddPCR. **(D)** Hybrid capture NGS with single-sided capture probes detected 52 sites with off-target integration in K562*attB* cells and most validated off-target frequencies were at or below our lower limit of detection. **(E)** Quantification of off-target insertion at CAS421 in K562*attB* cells using ddPCR confirms the frequency of off-target integration detected by hybrid capture NGS. **(F)** Table of 5 cryptic *attB* sites validated by WGS. Cryptic-seq discovered all 5 WGS-validated off-target sites but hybrid capture captured only 4 of the 5 sites detected by WGS. Cryptic-seq discovered all cryptic *attB* sites validated by WGS, HIDE-seq discovered only 1 of the WGS-validated sites. **(G)** Discovery assay classification errors for HIDE-seq and Cryptic-Seq reveal high level of false positive rate for Cryptic-seq but achieve the target goal of 0% false negative rate.

To qualify hybrid capture NGS for the quantitative detection of cryptic integration events we performed a standard curve experiment using commercially available purified human gDNA spiked with a custom synthetic DNA fragment at varying relative copy ratios. The DNA sequence of the fragments was designed to simulate integration at a HIDE-seq and Cryptic-seq discovered cryptic *attB* site CAS031 (chr6: 142247516- 142247561; GT dinucleotide located in minus strand starting at 142247538; **Figure 2c**). Notably, CAS031 was also discovered in a recent report of Bxb1 integrase off-target sites^18^. The synthetic DNA fragment was titrated into human gDNA at specific copy number concentrations (95%, 50%, 10%, 1%, 0.1%, 0.05%, 0.001%, and 0%) to generate an 8-point standard curve. These DNA standards were processed in triplicate by NGS library prep and target enrichment with hybrid capture panels containing probes hybridizing either left or right of CAS031. Illumina short-read sequencing and bioinformatic analysis was performed via our custom hybrid capture analysis pipeline. As shown in **Figure 4b**, by comparing observed versus expected integration frequencies, hybrid capture NGS affords accurate quantitation across standard curve and reliable detection of recombined reads (as defined as detection of the edit in ≥ 2/3 replicates) down to 0.1% and single-point detection with both left and right CAS031 probes and single point detection as low as 0.05% with the left probe at the level of sequencing performed in this experiment. For positive calling of integration events in hybrid capture NGS, we set the threshold that a read (UMI count) must be informative, i.e. the read spans the complete cryptic *attB* target sequence (unedited read) or the complete cryptic *attL* or *attR* sequence with a full P or P’ half-site sequence (integrated read). In this experiment, we recovered an average of 440 informative UMI counts for the left CAS031 probe samples and an average of 797 informative UMI counts for the right CAS031 probe samples (data not shown). We also observed strong correlation between observed and expected integration frequencies with Pearson r-squared values of 0.9897 and 0.9954 for the left and right CAS031 probes respectively, highlighting the quantitative ability of hybrid capture NGS for quantitative detection of cryptic integration events in edited gDNA (**Figure 4b**).

### Hybrid Capture NGS Validation of LSI off-target editing at cryptic *attB* sites in K562 cells

To examine the potential of HIDE-seq and Cryptic-seq to discover *bona fide* recombinogenic cryptic sites in cells, we leveraged an engineered K562 cell line that contains two *attB*-GT sequences in different genomic locations. This line was generated by transducing K562 cells with a lentivirus carrying insert in a PGK-*attB*-EF1α-PuroR cassette that integrated at unique genomic sites in both chromosome 6 and 17 (K562*attB* cells). Biological triplicate samples (Rep 1, Rep 2, and Rep 3) of K562*attB* cells were co-transfected with mRNA expressing Bxb1 integrase and an *attP*-GT plasmid donor, along with control samples in which K562*attB* cells were transfected with only the *attP*-GT plasmid donor (Control 1, Control 2, and Control 3), followed by harvesting and lysis 3 days post-transfection to isolate gDNA. Assessment of on-target editing by ddPCR showed robust on-target average integration frequencies of 45.5%, 48.4%, and 48.3% across edited Rep 1, Rep 2, and Rep 3 samples, respectively (**Figure 4c).**

To create a hybrid capture panel composed of discovered sites to test for off-target editing in K562s, we prioritized inclusion of cryptic sites located in Tier I (gene coding regions), Tier II (non-coding regions (exonic UTR or intronic) of coding genes, and Tier III (exonic or intronic regions of non-coding genes) as defined by the off-target threat matrix from The Broad Institute^35^ in combination with a prioritization of cryptic sites with UMI counts ≥ 10 from Cryptic-seq (**Figure 3c**). Of note, the probe panel consists of discovered cryptic sites and does not contain an on-target probe targeting the lentiviral cassette that contains the canonical *attB* sequence and, given the concordance in detection by left and right probes shown in **Figure 4b**, we designed the panel to contain only a single capture probe (either left or right) for each target (i.e., single sided panels). This inclusion criteria resulted in a single-sided probe panel size of 2,455 genomic sites. Of the 2,455 included sites, only 16 were discovered in HIDE-seq (CAS002, CAS005, CAS006, CAS008, CAS011, CAS013, CAS017, CAS018, CAS019, CAS020, CAS023, CAS024, CAS026, CAS030, CAS031, CAS032 in **Figure 2D**). For hybrid capture NGS, 2 of the edited K562*attB* biological replicates (Rep 1 and Rep 2) and 2 of the cargo alone samples (Control 1 and Control 2) were chosen for captured enrichment and analysis. Samples were sequenced on an Illumina NextSeq 2000 with 2x300 paired end sequencing, yielding an average of 3.7 million reads near probe across samples and an average target capture percentage of 37% (derived from quantifying reads near probes divided by total reads in the sample). FASTQ files were analyzed through a custom hybrid capture bioinformatic pipeline for indel and cryptic integration event detection.

For quantification of indels, we filtered targets containing ≥1,000 informative UMI average read counts across edited Rep 1 and Rep 2 and unedited Control 1 and Control 2 samples (to reduce any miscalls due to sampling bias). For a positive indel call, the indel must overlap within a 6 bp window surrounding the central GT dinucleotide with the minimal mapping criteria that the read that contains minimum of 50 bp of flanking each side of the dinucleotide. Using these inclusion criteria, 621 unique targets were analyzed for indel detection by comparing edited and unedited samples^33^. Statistical testing by Welch’s one-sided T test, revealed only 16 sites with p-values <0.05. However, across all 621 sites assayed, including the 16 sites with p-values <0.05, we identified only one potential off-target site, CAS14693, which was not significant but did display an Δindel frequency between the edited and control samples greater than 0.2% (data not shown). CAS14693 displayed averaged indel frequencies of 7.5% in edited samples and 6.7% in unedited samples with a Δindel frequency of ∼0.7% (data not shown). Visual inspection of the indel sequences indicates they are likely somatic indels present in the K562*attB* cell line as they represent precise and single CAG repeat deletions present in both edited and unedited samples in a CAG-rich repetitive region of the potential cryptic site (data not shown). In conclusion, of the 621 sites that passed our indel analysis criteria, we identified no off-target indels scored that associated with Bxb1 integrase activity.

For quantification of cryptic integration, positive calls were made on any informative genomic reads tagged at the GT dinucleotide with a full P or P’ half-site sequence from the DNA cargo in the UMI deduplicated read. In total, 52 cryptic sites were validated for off-target insertion in K562*attB* cells (12 sites in Rep1 and 48 sites in Rep2) with 8 sites showing recurrent off-target insertion in both Rep1 and Rep2 samples (CAS421, CAS023, CAS6083, CAS4310, CAS024, CAS985, CAS4668, CAS008) with editing frequencies ranging from 4.26% (CAS421, Rep 2) to 0.049% (CAS3972, Rep 2) (**Figure 4d**). Of the 52 unique validated sites across Rep 1 and Rep 2, 5 of the 52 sites were discovered in HIDE-seq (CAS002, CAS008, CAS023, CAS024, CAS031) while all 52 validated sites were discovered in Cryptic-seq (**Figure 4d**). It is important to note that K562 cells display inherent chromosomal copy number variation^36^, so integration frequencies of off-target edits may not be directly comparable between targets. Across the 52 validated sites, the minimum number of informed reads was 694 UMI counts for CAS761 with 1 UMI count supporting integration (Rep 2) and the maximum number of informed reads was 2,177 UMI counts for CAS3785 with 2 UMI counts supporting integration (Rep 2) (data not shown). To corroborate our findings with an orthogonal approach, we assessed off-target integration at CAS421 by ddPCR in Rep 1 and Rep 2 gDNA samples. Analysis by ddPCR reveal the integration frequencies of 2.96% and 5.91% in Rep 1 and Rep 2 respectively (**Figure 4e**), closely matches the integration frequencies at CAS421 via hybrid capture NGS (1.29% in Rep 1 and 4.26% in Rep 2) (**Figure 4d**). These confirmatory results at a single site by ddPCR, along with the validation data in **Figure 4b**, demonstrate the sensitivity and scalability of hybrid capture NGS for LSI off-target frequency determination.

### The LSI off-target characterization framework consisting of HIDE-Seq, Cyptic-Seq and Hybrid capture NGS, is more sensitive than WGS alone

To benchmark our discovery and validation framework against unbiased whole genome sequencing, we isolated gDNA from Rep 1 and Rep 2 edited K562*attB* cells **(Figure 4C**) and performed WGS to an average genomic coverage of 40x. To identify any Bxb1 integrase-mediated integration events from WGS, fgsv^26^ was used to call and filter any split reads that contain genomic sequences and either P or P’ sequences of the cargo tagged at the GT dinucleotide. Across the two WGS samples, we identified a total of 5 unique off-target insertions at sites CAS031 (Rep 1), CAS071 (Rep 1), CAS421 (Rep 1, Rep 2), and CAS985 (Rep1, Rep2) and CAS1991 (Rep 1) (**Figure 4f**). In terms of detection of these sites with our framework, 1 WGS validated site (CAS031) was discovered with HIDE-seq while all 5 WGS validated sites were discovered with Cryptic-seq (**Figure 4f).** In terms of validation, hybrid capture validated 4 of the 5 WGS validated sites (CAS031, CAS421, CAS985, CAS1991) with only CAS071 not called in the hybrid capture dataset. In contrast, WGS at 40x coverage did not detect the remaining 48 sites that were validated in the hybrid capture datasets (**Figure 4d**). Taken together, our results from both hybrid capture NGS and WGS validate a composite total of 53 off-target sites across the edited K562*attB* samples.

Examining the classification error rates described in Gillmore *et al.,*^37^ for CRISPR- Cas9 off-target discovery in the context of off-target characterization and potential off-target editing of a genome editing therapeutic, a high rate of false positives is tolerated in order to mitigate the potential for false negatives to occur. HIDE-seq discovered 36 potential cryptic *attB* sites (**Figure 2e**) with 16 of these sites present in the hybrid capture panel and 5 of the 16 tested sites displaying off-target editing in K562*attB* cells. As a result, HIDE-seq displays a false positive rate of 68.8% (1-(5 validated sites ÷ 16 tested discovered sites) and false negative rate of 90.6% (1-(5 validated sites÷53 validated sites) (**Figure 4g**). Cryptic-seq discovered 44,311 unique cryptic *attBs* with 2,455 of these sites present in the hybrid capture panel and 53 of these sites displaying off-target editing in K562*attB* cells. Accordingly, Cryptic-seq displays a false positive rate of 97.8% (1-(53 validated sites ÷ 2,455 tested discovered sites) and false negative rate of 0% (1-(53 validated sites ÷ 53 validated sites) (**Figure 4g**).

## DISCUSSION

Large serine integrases are a unique class of compact enzymes that have evolved to insert large DNA sequences (>50 kb) in specific genomic locations, and can catalyze all steps of the integration reaction, independent of host cell DNA repair pathways^2^. Thus, integrases have immense therapeutic potential genome editing^11-14, 18^. In particular, LSIs can be used in combination with Cas9 nickases and writing enzymes (e.g. reverse transcriptase^11, 14^ or ligases^38^) to enable therapeutic applications including endogenous gene replacement and the development of highly engineered cell-based medicines. LSIs allow for the directional, seamless integration of large DNA sequences without depending on FDEs or cellular DNA repair pathways. To date, comprehensive methods for detecting and validating LSI specificity have not been developed but will be required to develop integrase-based therapeutics.

Standard genotoxicity evaluation approaches, like the Ames test or Comet assays^39^, are not suitable for homology dependent off-target editing from an LSI or an RNA-guided nuclease because they are designed to detect the effects of homology independent sources DNA damage, such as chemical or physical agents. Homology dependent off-target editing poises the risk of recurrent gene disruption of a tumor suppressor that would be undetectable with standard approaches. Therefore, we have sought to contribute fit for purpose and sensitive methods to evaluate the specificity any LSI in the human genome to help advance the therapeutic application of endogenous gene replacement.

In contrast to iPSC-based cell therapy applications, where a clonal cell line is genome engineered and WGS alone is often sufficient to characterize off-target editing^40^, *ex-vivo* approaches that employ gene editing of bulk T cells or CD34+ hematopoietic stem and progenitor cells, as well as *in vivo* editing approaches that target the liver, require a two-step approach to first discover potential off-target sites and then validate sites with off-target edits through the detection off-target activity in edited cells^34, 37^.

The approaches we advance here comprise a comprehensive and sensitive suite of assays to discover and validate LSI off-target events to directly address regulatory and genotoxicity considerations for LSI-based strategies in gene and cell therapy approaches. We demonstrate that HIDE-seq is a readily applicable and unbiased biochemical-based WGS approach for identification of potential LSI off-target outcomes such as indels, cryptic integrations, and large structural variants. We increase the sensitivity of off-target discovery of cryptic integration events with Cryptic-seq, an ultra-sensitive biochemical approach that is greater than 1,200-fold more sensitive than HIDE-seq but is limited to detection of cryptic integrations. Cryptic-seq uses supraphysiological amounts of integrase and DNA template, significantly higher than a human cell would ever be exposed to in a therapeutic setting, followed by PCR enrichment, to identify a deep pool of off target sites for expansive analysis. We have demonstrated that this approach is extremely sensitive and has been specifically tuned to give a high false positive rate (>97%), increasing confidence that the total spectrum of potential off target sites with any integrase-dinucleotide combination will be discovered. Both HIDE-seq and Cryptic-seq can be adapted to accommodate multiple LSIs (i.e., multiplexing) or novelly designed LSIs. Importantly, both assays can be performed with genomic DNA isolated from therapeutically relevant cell types (e.g., human donor derived T cells, CD34+ hematopoietic stem and progenitor cells, primary human hepatocytes) or mixed populations of genomic DNA for more genetically relevant discovery of potential off-target sites. To readily perform scalable off-target validation of up to 1000s of potential LSI off-target sites in relevant edited cell types, we present a customized application of hybrid capture NGS that is tailored to accurately detect and quantify LSI off-target events, such as indels and cryptic integrations, which we benchmark to bulk WGS. In the context of the experimental data presented here, we demonstrate hybrid capture NGS has greater than 98% apparent validation hit rate (52/53 validated sites detected) compared to an apparent hit rate of WGS (at 40x coverage) of 9.4% (5/53 validated sites detected). By sharing our advances in molecular analytics for LSIs, which comprise powerful and promising tools for basic and clinical applications of programmable genomic insertion, we hope to help further the establishment of benchmarks for sensitivity of off-target detection in therapeutic paradigms that employ these fascinating enzymes.

## METHODS

### HEK293*attB* cell line generation

In-house produced lentivirus containing a transfer plasmid with an EF1α-PuroR-WPRE backbone containing a 46 bp Bxb1 *attB* insert (PL312) was transduced into HEK293 cells. Cells with Low MOI were plated in sterile 96 well plates under puromycin selection via serial dilutions for clone selection. Following clone selection, confirmation of lentiviral copy and identification of the lentivirus insertion site was performed using ligation mediated PCR with primers targeting the 5’ and 3’ LTRs along with Cergentis TLA mapping confirmed single lentiviral insertion on chromosome 5 (data not shown). Assessment of the insertion site by long read PCR with flanking genomic primers followed by Oxford Nanopore Technologies long read sequencing confirmed the lentiviral insert sequence, which had partially concatemerized to ultimately contain two Bxb1 *attB* sequences, separated by 4,272 bp of intervening lentiviral sequence.

### K562*attB* cell line generation

Lentiviral plasmid (PL2811) with a PGK-EF1α-PuroR backbone containing a 46 bp Bxb1 *attB* inserts was acquired from GenScript and used for lentiviral production at Azenta. To generate engineered cell lines containing *attB* sequences, K562 cells (ATCC, Cat# CCL- 243) were transduced with lentivirus doses of 5 µL, 10 µL, 20 µL, 30 µL and 50 µL infected by spinfection at 1000g for 30 minutes at 33°C to find appropriate dose of lentivirus with low multiplicity of infection (MOI). Cells with Low MOI were plated in sterile 96 well plates under puromycin selection via serial dilutions for clone selection. Following clone selection, confirmation of lentivirus insertion site was performed using Lenti-X™ Integration Site Analysis Kit (Takara) and WGS, which both confirmed the presence of single lentiviral integration events on chromosome 6 and 17.

### HIDE-seq

HIDE-seq reactions with 8 µg of purified gDNA from PBMCs (Qiagen Blood and Tissue kit) or, in separate reactions, 8 µg of purified gDNA HEK293*attB* cells was incubated with 10 nM attB_66bp annealed oligonucleotide (for cryptic *attP* site or FDE discovery) or 10 nM attP_72bp annealed oligonucleotide (for cryptic *attB* site or FDE discovery) and 400 nM of recombinant Bxb1 integrase (GenScript) in recombination buffer^7^ (10 nM Tris-HCl, pH 8.0, 100 mM KCl, 5% glycerol) for 8 hours at 37°C and then held at 4°C. To anneal the attB_66 and attP_72 oligonucleotide substrates for HIDE-seq, 25 µl of attB_66bp_P_Top (100 mM) and 25 µl of attB_66bp_P_Bot (100 mM) or 25 µl of attP_72bp_P_Top (100 mM) and 25 µl of attP_72bp_P_Bot (100 mM), respectively were mixed together in wells of a 20 µl 96 well plate. Note, oligonucleotides were resuspended in duplex buffer (Integrated DNA technologies (IDT)). The plate with oligonucleotide samples were placed into a thermocycler, and the following anneal program was performed with a 105°C heated lid: 95°C/2 minutes, 750 cycles at -0.1°C per cycle/1s, 4°C hold. Reactions were inactivated with RNAase A (New England Biolabs (NEB) Monarch) and then Proteinase K (NEB) and gDNA was purified with 0.7x AMPure (Beckmann Coulter) bead cleanup. Control Digenome-seq reactions were performed with gDNA and spCas9 (IDT) with VEGFA S2 sgRNA (IDT) were performed according to the published protocol^21^. Isolated gDNA from both HIDE-seq and Digenome-seq reactions was quantified by Qubit fluorometry (Thermo Fisher Scientific) and submitted for PCR-free WGS at 30x coverage (Azenta). FASTQ files from Illumina sequencing were loaded into a custom bioinformatic pipeline for HIDE-seq called tbDigIn (https://github.com/didacs/tbDigIn) developed by Tome Biosciences and Fulcrum Genomics which quantified the number of unclipped reads (for FDE detection) or soft-clipped reads (recombined sites). The bioinformatic workflow for tbDigIn starts with the alignment of sequencing reads from FASTQ files against the hg38 human reference genome^41^ with appended *attP* and *attB* sequences by BWA aligner^42^ to create mapped BAM files for each sample. The resulting BAM files are deduplicated by Picard^43^ and inputted into HIDE, a modified Digenomitas^24^ pipeline to output .csv files containing sites per sample as clipped counts for integration events and unclipped counts for FDEs.

### Cryptic-seq

HEK293*attB* gDNA was incubated with Tn5 transposase^27^ (custom purification by GenScript) pre-annealed with MES_Rev_3InvdR and MES_AmpSeq_P5 oligos) for 7 minutes at 55°C to tagment the gDNA and shear it to an average size of approximately 700bp. The Cryptic-seq donor plasmid (PL2312) includes an *attP* integrase attachment site with a GT dinucleotide flanked by a unique 12 bp barcode sequence (BC1) and Illumina N7 primer binding site (PBS) on the 5’ side of the *attP* and a unique barcode sequence (BC2) sequence on and an Illumina P7 sequencing primer binding site (PBS) on the 3’ side of the *attP*. The biochemical reaction was performed with 1 µg tagmented gDNA, 10 nM of the Cryptic-seq donor plasmid, and 1 µM integrase and incubating these components at 37°C for 4h in a recombination buffer^7^. The reaction was stopped by adding sodium dodecyl sulfate to a final concentration of 0.1% and the products were cleaned using the Zymo clean and concentrator kit (Zymo research).

PCR was used to amplify the regions of integration using Q5 polymerase (NEB) and primers for P5 containing distinct barcodes for each sample (LM_P5_F1-F3) and a single N7 primer (GN037_CrypticSeq_N7_i7_N701). Following PCR, a 1.5x AMPure bead clean-up was performed, and products were run on a Tapestation 4200 (Agilent Technologies) using D1000 tape to confirm amplification. Library concentration was determined using the Next Library Quant Kit (NEB) for Illumina. Libraries were normalized to 2nM, loaded at a concentration of 750pM and sequenced via Illumina Next Seq according to the manufacturer’s instructions.

For bioinformatic analysis of cryptic-seq data, FASTQ files from Illumina sequencing were loaded into a custom cryptic-seq bioinformatic pipeline tbChaSIn (https://github.com/didacs/tbChaSIn) developed by Tome Biosciences and Fulcrum Genomics to discover and quantify cryptic recombination sites from Cryptic-seq data. The bioinformatic workflow starts with the trimming of reads that contain leading *attP* or *attB* sequences (P, P’ or B, B’) followed by a search and trim of any Tn5 mosaic end (ME) sequence (CTGTCTCTTATACACATCT, Illumina) in the reads. Reads trimmed for att and ME sequences are aligned against the hg38 human reference genome^41^ with appended *attP* and *attB* sequences by BWA aligner^42^ to create mapped BAM files for each sample. The resulting BAM files are deduplicated by Picard^43^ and queried for integration sites with to generate output .csv files containing sites per sample along with collation of sites across samples to generate output .csv files containing collated sites.

### Hybrid Capture Quantitative Validation with DNA standards

Simulated LSR-mediated cargo integration for CAS031 DNA fragments were synthesized as gBlocks (IDT). gBlocks were designed to include approximately 200 bp left and right of the recombination junction (*attL* or *attR*) dinucleotide followed by approximately 1300 bp of random DNA stuffer sequence at each end to achieve a fragment length of approximately 3000 bp for optimal DNA fragmentation (CHR6_RC_Off_PL753_attL, CHR6_RC_Off_PL753_attR). Standard curve titrations were prepared by spiking in gBlocks into human gDNA (Promega catalog #G304A) at molar ratios representing 95%, 50%, 10%, 1%, 0.1%, 0.05%, 0.001%, and 0% I-PGI. Samples were fragmented to an average size of 550 bp via sonication (Covaris ME220) and NGS libraries prepared according to the IDT xGen DNA Library Prep Kit MC UNI (Version 2) protocol using xGen™ UDI-UMI Adapters (IDT catalog #10005903). Target enrichment was performed according to xGen™ hybridization capture of DNA libraries (Version 7) protocol using custom hybrid capture panels containing probes either left (5’) or right (3’) of CAS031. Hybrid capture libraries were sequenced by Illumina short-read sequencing via NextSeq 2000 paired-end (2 x 150 bp) sequencing using standard NextSeq 2000 P3 Reagents (300-Cycles) (Illumina catalog #20040561) and analyzed using a custom hybrid capture bioinformatics pipeline called tbREVEAL developed at Tome Biosciences (described below).

### Editing of K562*attB* cell lines

K562*attB* cells were maintained in RMPI1640 media (Gibco) +10% fetal bovine serum (Gibco) + 2ug/ml puromycin (Life Technologies) and passaged at ratio of 0.1 million cells per ml every 3-5 days. For editing, 200,000 cells were electroporated with 3 µg mRNA for Bxb1 integrase and 3 µg of a donor construct containing *attP* integrase attachment sites and a GT central dinucleotide using the Lonza 4D-Nucleofector™ X Unit according to the manufacturer’s instructions for K562 cells. After 3 days, 80% of the cells were collected for gDNA extraction using the Qiagen blood and tissue kit and the remaining 20% were expanded then banked.

### Hybrid capture NGS of edited K562*attB* cells

Genomic DNA extracted from edited K562*attB* cells were used as input for hybridization capture NGS. A Covaris ME220 focused ultrasonicator was used to shear 400 ng of Rep 1, Rep 2, Control 1, and Control 2 gDNA samples to an average size of 400 bp. Fragmented DNA was end repaired and A-tailed followed by ligation of Illumina sequencing adapters using Twist library preparation kit (cat#104177) to create Illumina paired end sequencing libraries. Hybridization, capture and post capture amplification of the prepared libraries to probes was performed using Twist target enrichment standard hybridization kit as per manufacturer’s instructions (Twist Library Preparation Kit 1, Mechanical Fragmentation, cat # 100876). Libraries were sequenced on Illumina NextSeq 2000 platform using a P2 600 cycle kit (Illumina) according to standard manufacturer’s protocol.

For bioinformatic analysis of hybrid capture NGS data, FASTQ files from Illumina sequencing were loaded into a custom bioinformatic pipeline called tbREVEAL (https://github.com/jessie-wangjie/tbREVEAL) developed by Tome Biosciences to quantify edited reads corresponding to unedited reads, reads containing indels, reads containing cargo DNA sequence and thus represent recombined reads. Briefly, input data in FASTQ or BAM format undergo quality control using FASTP (https://github.com/OpenGene/fastp), which includes adapter trimming, quality filtering, and extraction of unique molecular identifiers (UMIs). Additionally, paired-end reads are merged into a single file. The processed reads are aligned to the hg38 reference genome^41^ using BWA aligner^42^. UMIs are deduplicated from the aligned BAM files to remove PCR duplicates. Target information is collected and used to generate reference amplicons. Target reads are extracted from the deduplicated BAM files and aligned to the reference amplicons using BWA aligner^42^. Editing events are quantified using a custom Python script designed to analyze the aligned target reads. Visualization of editing events is performed using a custom script to generate graphical representations. Editing sites are collated across samples are outputted into an excel spreadsheet using a custom script. HTML reports by papermill (https://papermill.readthedocs.io/en/latest/index.html) are generated to summarize the results of the analysis.

### Whole genome sequencing of edited *K562attB* cells

For WGS, Bxb1 integrase-edited Rep 1 and Rep 2 K562*attB* gDNA genomic DNA was extracted using DNeasy blood and tissue kit (Qiagen) and samples library preparation and Illumina short read sequencing with a target of 60x genomic coverage. For library preparation, genomic DNA was sheared using a Covaris ME220 sonicator to average size of 400 bp, sheared fragments were end repaired, A-tailed and ligated with Illumina sequencing adapters and the resulting library was sequenced on an Illumina NovaSeq platform (Azenta). For bioinformatic analysis of WGS data, FASTQ files from Illumina sequencing were mapped to the hg38 reference genome and cargo reference using BWA^42^ followed by fgsv^26^ from Fulcrum Genomics was used to predict potential chimeric reads. Chimeric reads with one breakpoint at the reference genome and with the second breakpoint at the *attP* di-nucleotides in the DNA cargo reference were retained for further investigation.

## MATERIALS

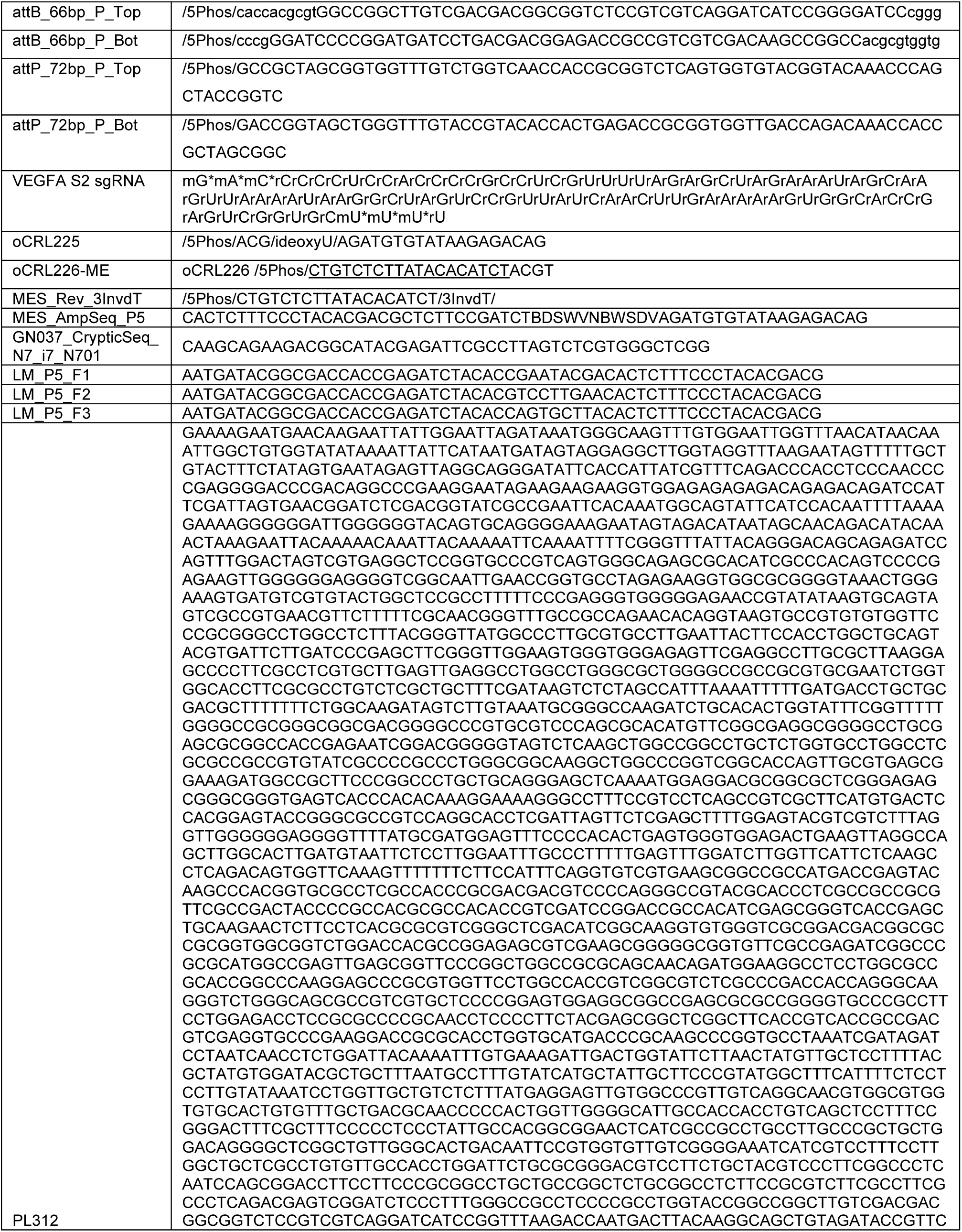
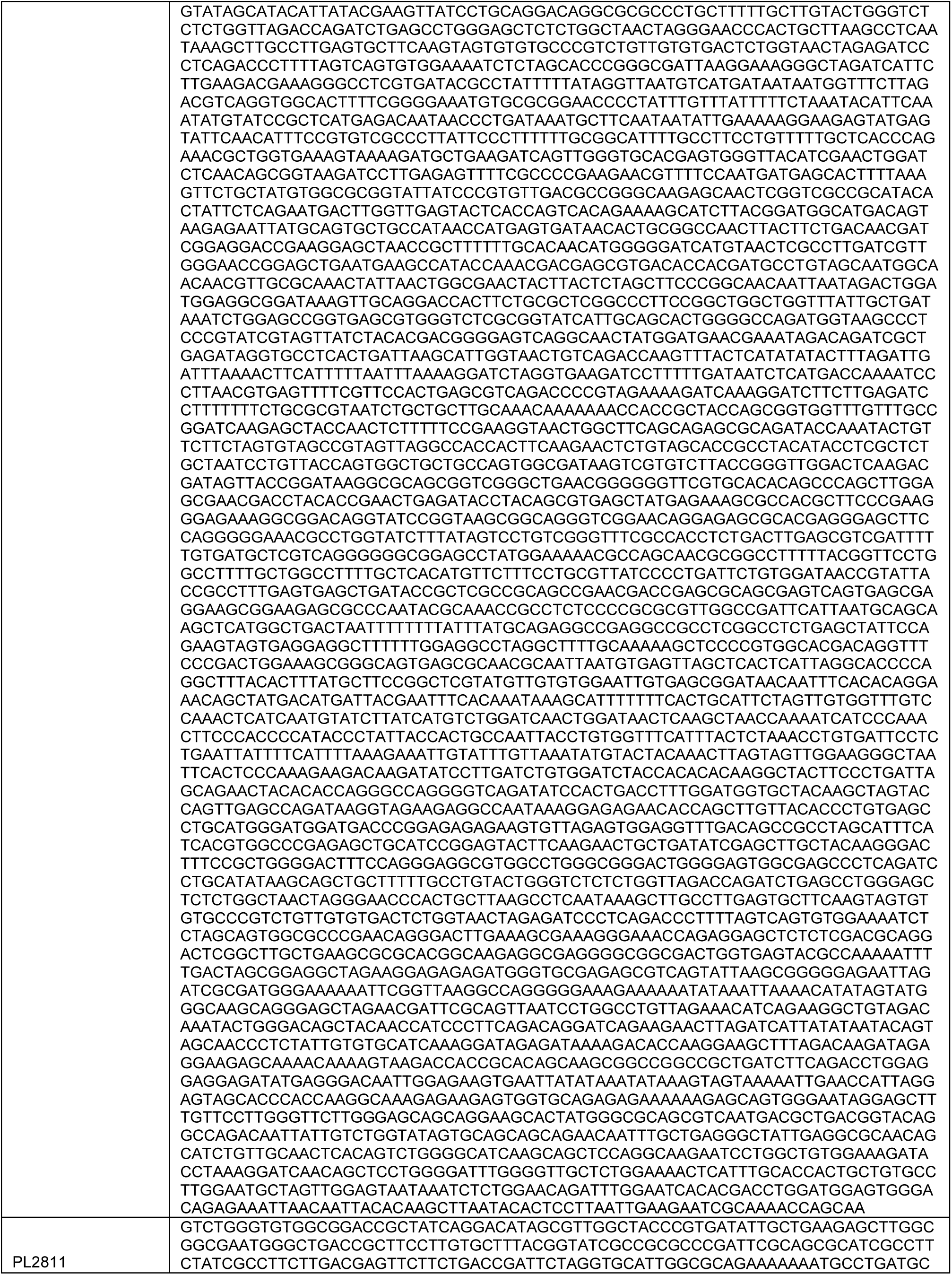
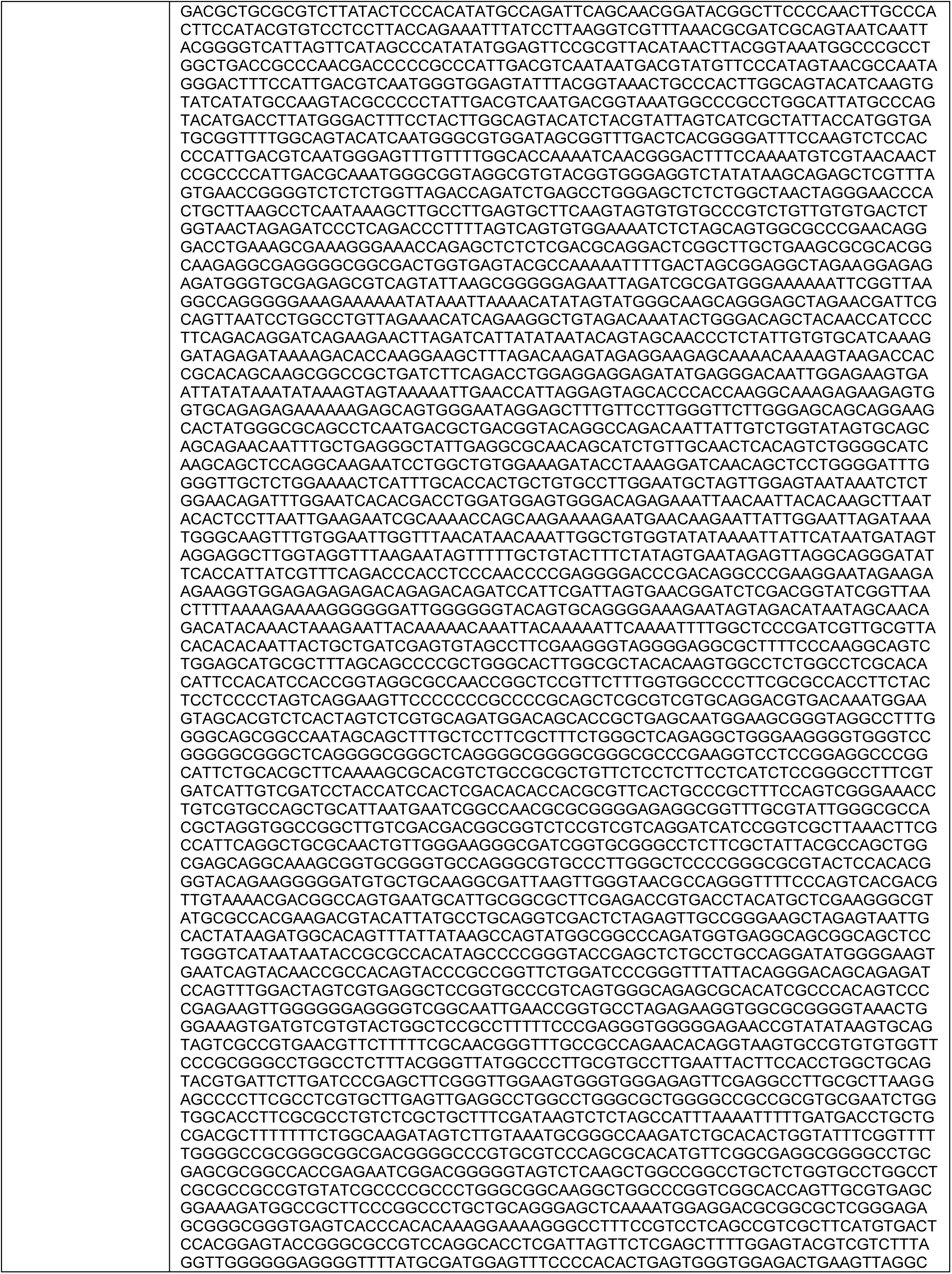
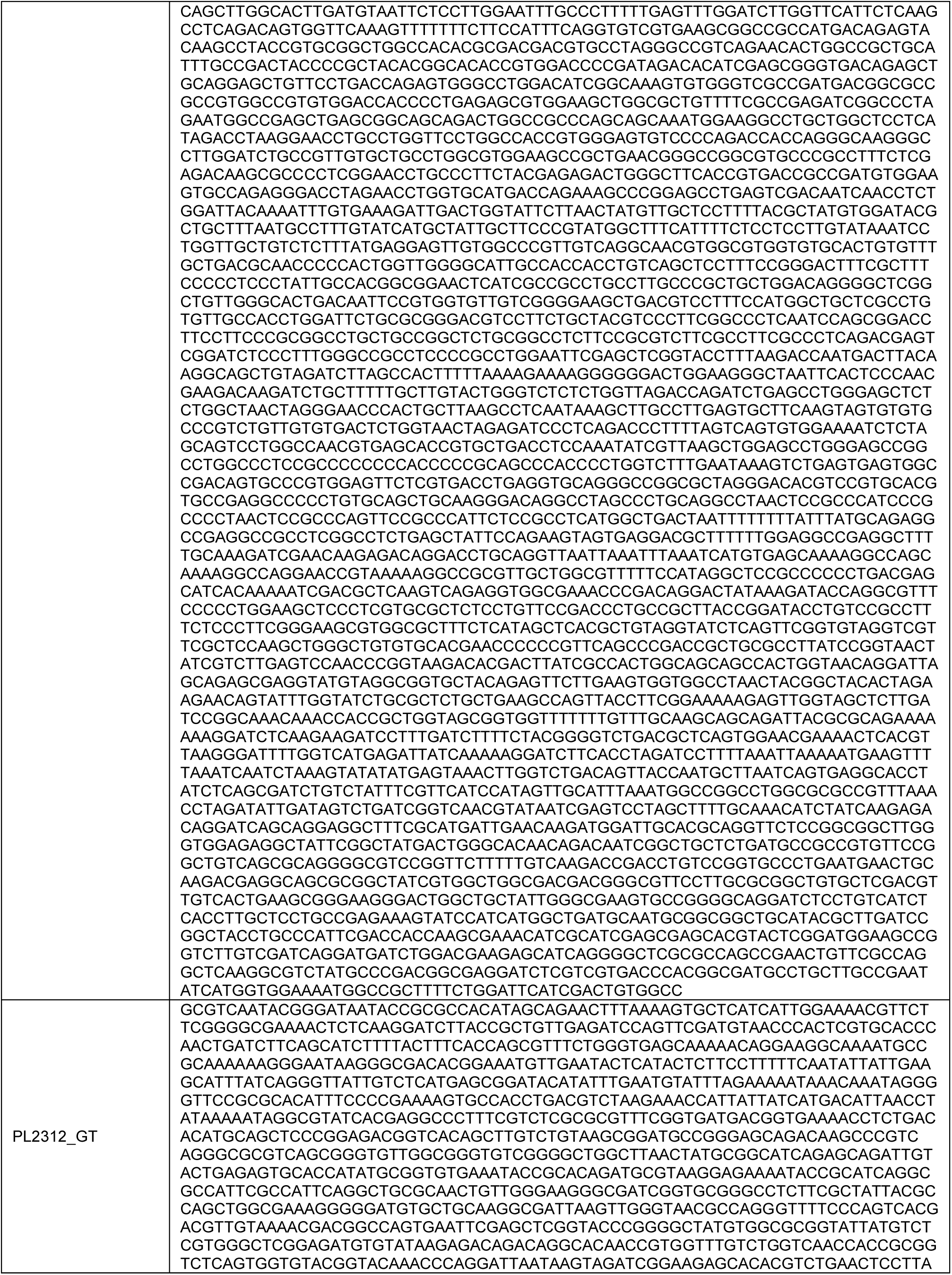
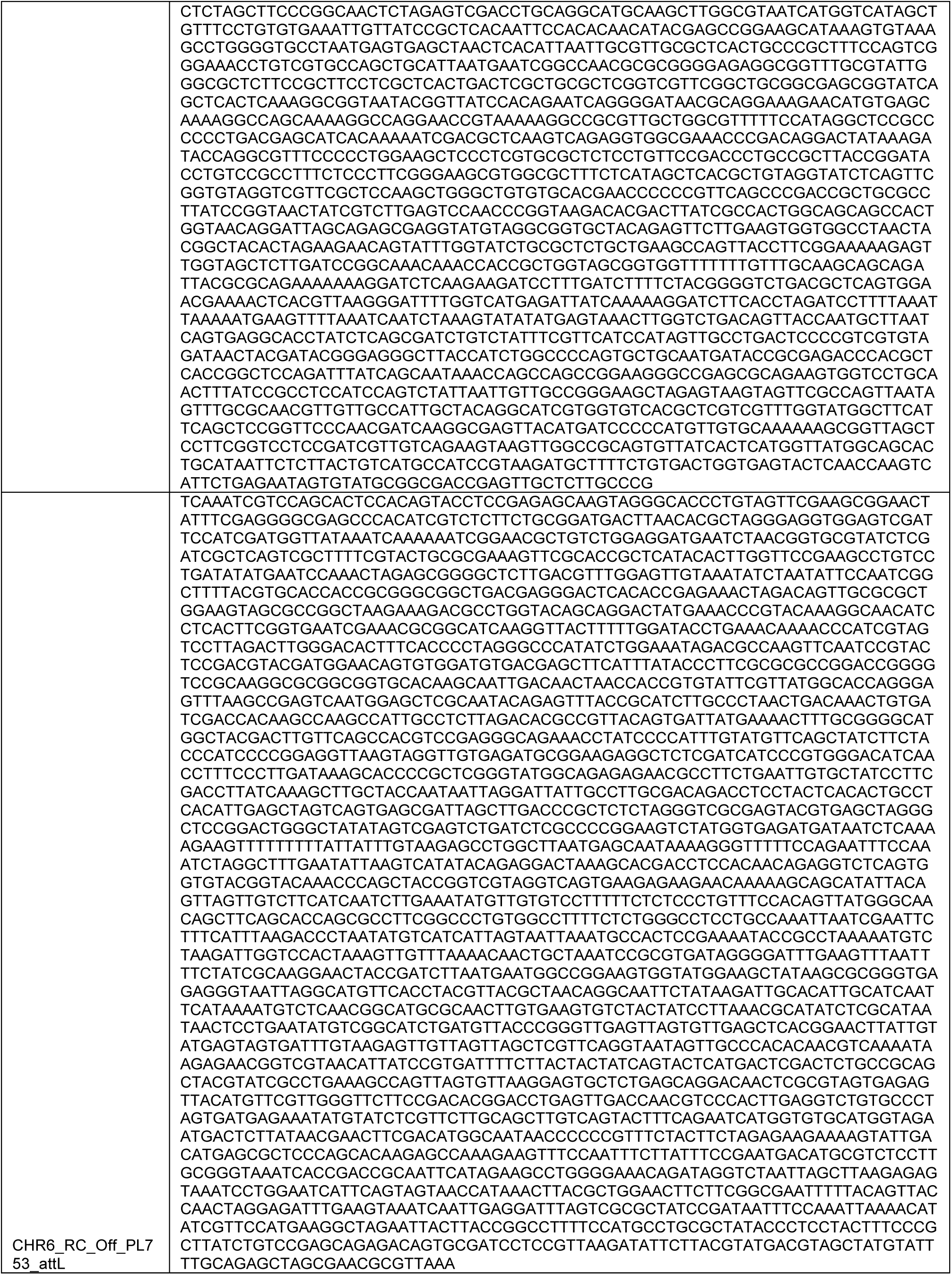
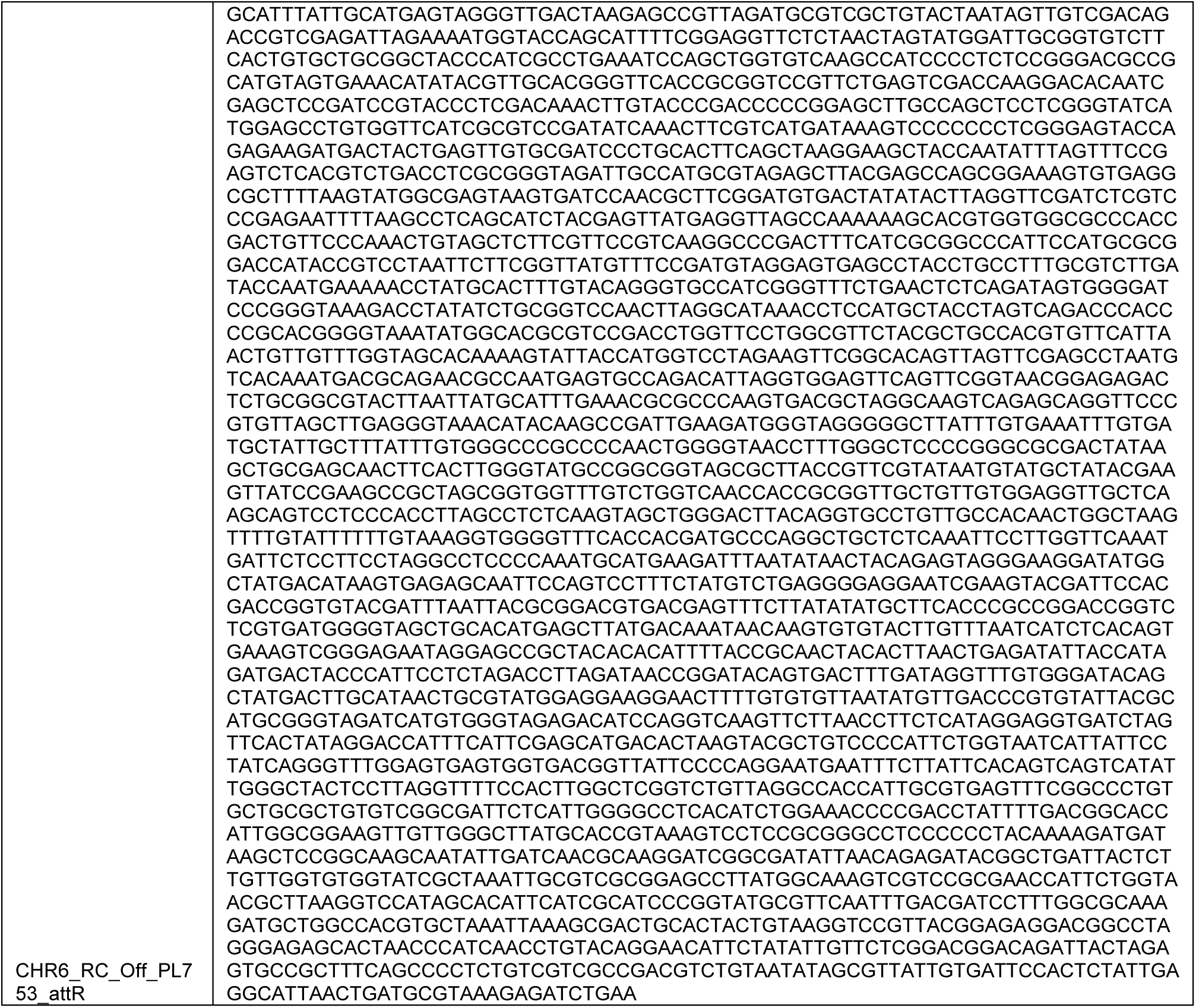

## DATA AND SOFTWARE AVAILABILITY

The HIDE-seq, Cryptic-seq, and Hybrid Capture analysis pipelines are accessible through GitHub at https://github.com/didacs/tbDigIn, https://github.com/didacs/tbChaSIn, https://github.com/jessie-wangjie/tbREVEAL, respectively. The project number for the next-generation sequencing data reported in this paper is NCBI SRA: <u>PRJNA1167015</u>. Genomic coordinates (hg38) of HIDE-seq and Cryptic-seq discovered sites are detailed in Supplementary Tables 1 and 2, respectively.

## ACKNOWLEDGEMENTS

We acknowledge all members of Tome Biosciences for their support in the generation of this work. Figures 1B-C, 2A, 3A, and 4A were created with Biorender.com.

